# Crosslinkers both drive and brake cytoskeletal remodeling and furrowing in cytokinesis

**DOI:** 10.1101/150813

**Authors:** Carlos Patino Descovich, Daniel B. Cortes, Sean Ryan, Jazmine Nash, Li Zhang, Paul S. Maddox, Francois Nedelec, Amy Shaub Maddox

## Abstract

Cytokinesis and other cell shape changes are driven by the actomyosin contractile cytoskeleton. The molecular rearrangements that bring about contractility in non-muscle cells are currently debated. Specifically, both filament sliding by myosin motors, as well as cytoskeletal crosslinking by myosins and non-motor crosslinkers, are thought to promote contractility. Here, we examined how the abundance of motor and non-motor crosslinkers controls the speed of cytokinetic furrowing. We built a minimal model to simulate the contractile dynamics of the *C. elegans* zygote cytokinetic ring. This model predicted that intermediate levels of non-motor crosslinkers would allow maximal contraction speed, which we found to be the case for the scaffold protein anillin, *in vivo*. Our model also demonstrated a non-linear relationship between the abundance of motor ensembles and contraction speed. *In vivo*, thorough depletion of non-muscle myosin II delayed furrow initiation, slowed F-actin alignment, and reduced maximum contraction speed, but partial depletion allowed faster-than-expected kinetics. Thus, both motor and non-motor crosslinkers promote cytokinetic ring closure when present at low levels, but act as a brake when present at higher levels. Together, our findings extend the growing appreciation for the roles of crosslinkers, but reveal that they not only drive but also brake cytoskeletal remodeling.

## Introduction

The actomyosin cortex powers cell shape change during diverse cellular behaviors including cell migration, tissue morphogenesis and cell division. The cortex is a heterogeneous meshwork rich in actin filaments (F-actin), cytoskeletal crosslinkers, cytoskeleton-plasma membrane linkers, and myosin motors (Clark *et al.*, 2014). The final step of mitotic or meiotic cell division is cytokinesis, the physical division of the cell into two. During cytokinesis, cues from the anaphase spindle elicit specialization of the cortical cytoskeleton to form an actomyosin ring at the cell equator. The cytokinetic actomyosin ring is enriched with formin family actin nucleators that generate long unbranched F-actin, active non-muscle myosin II (NMM-II), and crosslinking proteins including anillin, alpha-actinin, the septins, and plastin. The ring dynamically rearranges, initially from a wide band into a tighter ring, and ultimately completely closes, generating a membrane partition between the daughter cells (Green *et al.*, 2012).

The cytoskeletal rearrangements proposed to bring about cytokinetic ring closure include actomyosin filament sliding, F-actin depolymerization, and cytoskeletal crosslinking (Mendes Pinto *et al.*, 2013). Sliding is thought to be driven by the motor activity of myosins including NMM-II, and loss of NMM-II function via injection of a blocking antibody, gene knock-out, protein depletion, or pharmaceutical inhibition eliminates furrowing (Schroeder, 1972; Mabuchi and Okuno, 1977; Straight *et al.*, 2003; Matsumura *et al.*, 2011). In some settings, the rate of cell constriction scales with the motor activity of NMM-II (Vasquez *et al.*, 2016). However, a motor-impaired NMM-II mutant that crosslinks, but does not slide, F-actin rescued the speed of cytokinetic furrowing caused by thorough depletion or genetic deletion of NMM-II from animal cells (Ma *et al.*, 2012). In addition, cell biological work with budding yeast and theoretical modeling demonstrated that ring closure can be powered by the thermal ratcheting of non-motor crosslinkers along depolymerizing F-actin (Mendes Pinto *et al.*, 2012; Oelz *et al.*, 2015). Similarly, in fission yeast, a motor-impaired NMM-II mutant that robustly crosslinks F-actin is sufficient for ring assembly (Palani *et al.*, 2017). Collectively, these studies suggest that F-actin crosslinking promotes cytoskeletal remodeling during cytokinesis, and ring closure in animal and fungal cells.

As introduced above, several cytoskeletal crosslinkers are abundant in the cytokinetic ring in animal cells, including NMM-II and anillin. NMM-II forms minifilaments, bipolar assemblies containing 16-56 motor subunits, each of which is bound to F-actin for 5-35% of its ATPase cycle (Burridge and Bray, 1975; Niederman and Pollard, 1975; Sinard *et al.*, 1989; Verkhovsky *et al.*, 1995). NMM-II minifilaments are robust F-actin crosslinkers (Vicente-Manzanares *et al.*, 2009). Anillin is a multi-domain protein thought to coordinate both structural and regulatory elements of the cytokinetic ring. Anillin was first identified as an F-actin bundling protein, and has since been shown to bind active NMM-II, septins, the formin mDia2, RhoA, and several other proteins, as well as plasma membrane lipids (Piekny and Maddox, 2010; Liu *et al.*, 2012; Reyes *et al.*, 2014; Sun *et al.*, 2015). Anillin depletion causes cytokinesis failure in cultured mammalian and *Drosophila* cells, but is dispensable for cytokinetic ring closure in the *C. elegans* zygote (Maddox *et al.*, 2005; Straight *et al.*, 2005; Zhao and Fang, 2005; Hickson and O’Farrell, 2008). Cortexillin crosslinks F-actin indirectly to the membrane and is important for the mechanical properties of the cortex and cytokinetic dynamics in the social amoeba *Dictyostelium discoideum* (Srivastava *et al.*, 2016). Additional cytoskeletal crosslinkers that constitutively reside in the cortex, such as fimbrin/plastin, also contribute to cytokinesis (Ding *et al.*, 2017). In sum, diverse cytoskeletal crosslinkers are abundant in the cytokinetic ring.

Interestingly, before crosslinking was implicated in driving cytokinesis, it was demonstrated that crosslinkers can block ring closure in mammalian cultured cells. The abundance of alpha-actinin scales inversely with furrowing speed in animal cells, as overexpression slows furrowing while depletion allows faster furrowing (Mukhina *et al.*, 2007). The picture is more complex in fission yeast, where an intermediate abundance and F-actin binding affinity of alpha-actinin are required for proper cytokinetic ring dynamics; thus this crosslinker both drives and blocks cytokinetic ring remodeling (Li *et al.*, 2016). Similarly, *in vitro* reconstituted and simulated simple actomyosin rings exhibit maximal contraction speed and force generation with an intermediate amount of connectivity (Ennomani *et al.*, 2016). Finally, an experimental increase or decrease of the abundance of plastin/fimbrin slows cytokinetic furrowing in the *C. elegans* zygote (Ding *et al.*, 2017). Thus, the abundance of crosslinkers has been variously shown to scale with actomyosin ring closure speed positively, negatively, or in a complex manner (speed maximal at intermediate crosslinker levels). As such, it is currently unresolved whether cytoskeletal crosslinkers drive or attenuate remodeling in the cytokinetic ring.

Here, we tested how tuning motor- and non-motor crosslinkers influences cytokinetic ring constriction in animal cells. We used the *C. elegans* zygote as a model cell type since its stereotyped size, shape, and cell division kinetics, and its mechanical isolation within the eggshell are well-suited for quantitative studies of ring-intrinsic factors. To develop intuition about tuning crosslinker levels in this contractile system and guide our biological experimentation, we built an agent-based minimal model of the *C. elegans* zygote cytokinetic ring, depicting fibers representing F-actin, fiber crosslinkers, and motor ensembles representing NMM-II minifilaments. Tuning crosslinker abundance *in silico* predicted that an intermediate level of non-motor crosslinker would allow maximal ring closure speed. We then targeted the scaffold protein anillin *in vivo*, and generated populations of cells with a graded abundance of anillin by carrying out RNA-mediated depletion over a time-course. Partial depletion of anillin allowed faster furrowing than observed in control cells, but in more thoroughly depleted cells, speed was normal. We next used our model to tune the abundance of motor ensembles and found a non-linear relationship between motor abundance and ring closure speed. *In vivo*, partial NMM-II depletion allowed faster furrowing than in control cells, while thorough depletion slowed ring closure. Towards defining the mechanism by which NMM-II both drives and brakes furrowing, we examined the kinetics of F-actin organization and found evidence that NMM-II also not only drives but also slows cytoskeletal remodeling. Our work demonstrates that both a motor- and a non-motor crosslinker can not only drive, but also attenuate, cytokinesis and thus extends our current understanding of the roles of cytoskeletal crosslinkers in cortical remodeling.

## Results

### Simulated actomyosin rings with NMM-II-like motor ensembles close with in vivo kinetics

Agent-based simulation of cytoskeletal machinery by CytoSim was recently used to tune the abundance of motors and non-motor crosslinkers in actomyosin-like patches and rings with different architectures (Ennomani *et al.*, 2016)While this approach provided the valuable insight that intermediate levels of crosslinkers allow maximal contraction speed, several features of the published models limit their relevance to cytokinesis *in vivo*. First, they employed motors modeled on myosin VI (a dimeric, pointed end-directed, highly processive motor (Rock *et al.*, 2001)), or on a generic, fast, processive barbed end-directed motor (Ding *et al.*, 2017). Furthermore, to model *in vitro* reconstituted rings with stabilized F-actin, simulated fibers were non-dynamic (Ennomani *et al.*, 2016). Here, we built on published models to depict NMM-II-like motors (see below), crosslinkers, and dynamic fibers, in a 30 micron-diameter ring representing a two-dimensional slice of the *C. elegans* zygote cytokinetic ring. The absolute and relative abundance of components was set according to measurements of the fission yeast cytokinetic ring, scaled to a cross section of the *C. elegans* zygote division plane (Figure 1A; (Wu and Pollard, 2005); see Methods). The starting length and treadmilling dynamics of actin fibers were based on measurements from molecule counting studies, electron microscopy, as well as previously-established parameter ranges (see Supplemental Table 1) (Wu and Pollard, 2005; Kamasaki *et al.*, 2007; Davies *et al.*, 2014).

**Figure 1.**
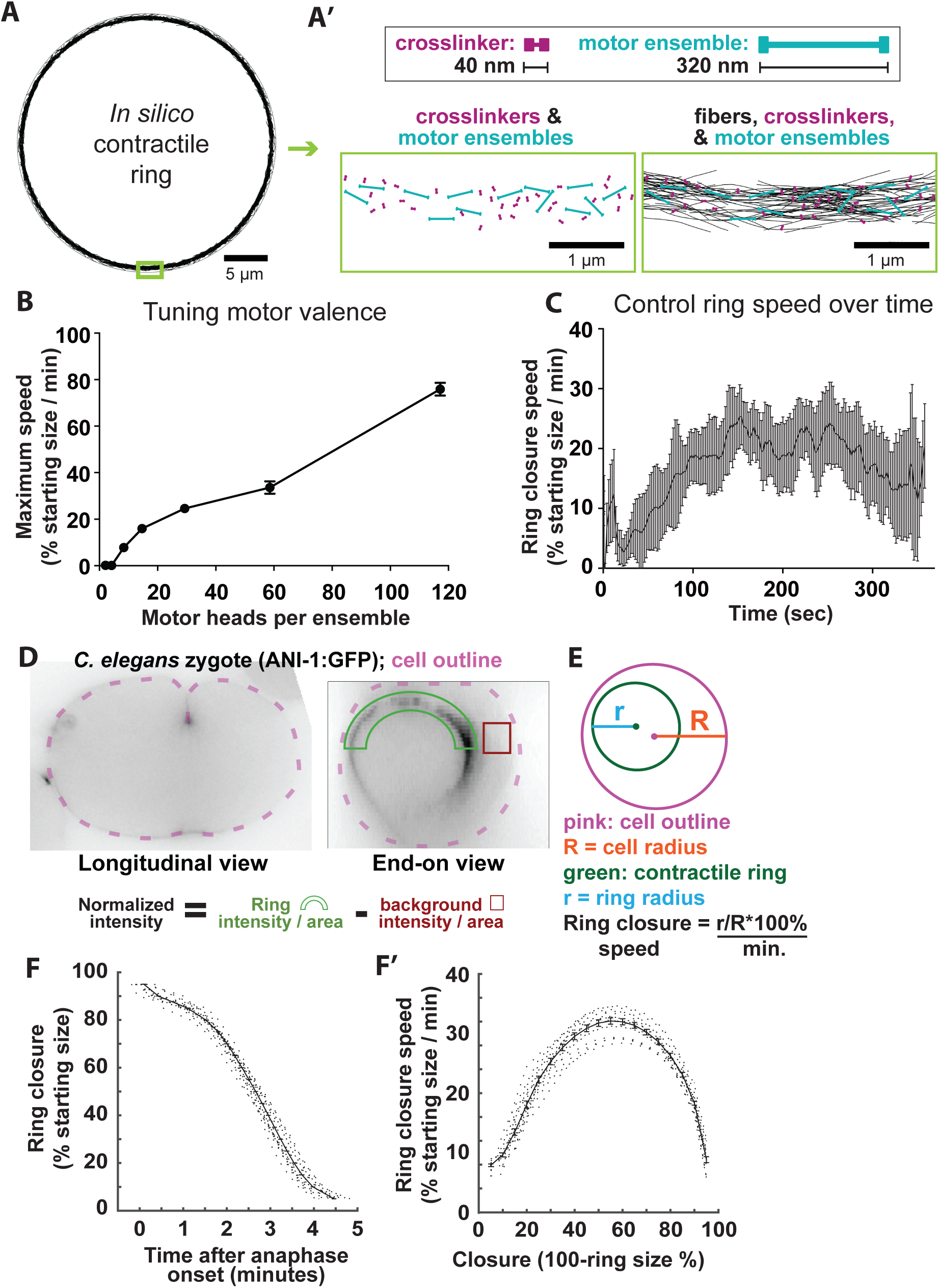
An intermediate abundance of non-motor crosslinkers allows maximal contractile ring closure speed in silico. A) In silico contractile ring at timepoint 0. A’) Top: scale drawings of crosslinkers and mono-ensembles. Below: magnified views of crosslinkers and motor ensembles without fibers, and all three components. B) Maximum ring closure speed for simulations varying the motor rod behavior to represent a range of subunits per motor ensemble (2 to 112 subunits per motor ensemble). Increasing motor ensemble valence increases maximum speed, but above 60 subunits, maximum speed was biologically irrelevant. nw = 4. C) Average ± SD closure speed of five control simulations. D) Transverse and end-on optical sections of C. elegans zygotes expressing fluorescently-tagged anillin (mNG::ANI-1). Green arc and red box represent regions measured for cytokinetic ring fluorescence intensity and background normalization, respectively. E) Schematic of ring size measurements for calculation of closure speed. F) Ring closure over time plots for control cells (n = 32). F’) Furrowing speed over time calculated from control cell data in F.

While employing many established parameters in our model, we adapted published models to simulate the behavior of NMM-II minifilaments, modeling motor ensembles as rods with a motoring fiber-binding site (“hand”) at each end. The behavior of each hand exhibits the collective binding dynamics for a prescribed number of motor heads according to alterations of motor parameters that include binding rate, unbinding rate, and duty ratio (see Supplemental Table 1) (Stam *et al.*, 2015). Since NMM-II can exist as dimers or as minifilaments with as many as 56 motor subunits *in vivo* (Burridge and Bray, 1975; Niederman and Pollard, 1975; Sinard et al., 1989; Verkhovsky et al., 1995), we kept the number of motor ensembles constant and tested how the speed of *in silico* ring closure was affected by the number of motor subunits per ensemble (the valence). Rings with motor ensembles representing 28 motor subunits closed with the most biologically relevant speeds (Figure 1B) leading us to utilize this valence in our control simulations.

We then measured the kinetics of simulated ring closure and found that our control *in silico* rings accelerated, maintained a relatively constant speed for much of closure, with maximum ring closure rates occurring 150-250 seconds after closure initiation, before gradually decelerating (Figure 1C). To compare simulation results with *in vivo* data, we acquired optical sections throughout the thickness of *C. elegans* zygotes, rotated image data sets 90 degrees to observe the entire division plane, visualized with a fluorescently-tagged ring component, and used custom Matlab-based software to annotate the position of the cytokinetic ring over time (Figure 1D, E; (Dorn *et al.*, 2010)). As *in silico*, the ring closed in an average of 5 minutes, accelerating until approximately half closed, and then decelerating (Figure 1F, F’; (Bourdages *et al.*, 2014)). Thus, simulated control rings and rings visualized *in vivo* exhibited similar closure dynamics, suggesting that our *in silico* approach using bipolar motor ensembles faithfully recapitulates some aspects of *in vivo* ring dynamics.

### An intermediate amount of non-motor crosslinkers allows maximal ring closure speed in silico and in vivo

As introduced above, previous work has demonstrated that intermediate levels of cytoskeletal crosslinking maximizes constriction of acto-myosin VI networks *in vitro* and *in silico*, and the fission yeast and *C. elegans* cytokinetic rings *in vivo* (Ennomani *et al.*, 2016; Li *et al.*, 2016; Ding *et al.*, 2017). To test the dependence of *in silico* ring constriction on non-motor fiber crosslinkers in our simulations, we varied the number of crosslinkers around the value set according to the measured collective abundance of two major F-actin crosslinkers (alpha-actinin and anillin (Mid1)) in the fission yeast cytokinetic ring (Wu and Pollard, 2005), scaled up to our 30-micron diameter ring. At the starting value (12,000) set from estimates of fission yeast, rings often fragmented but the maximum closure speed of those that remained intact was essentially unchanged (Figure 2A). Since ring fragmentation is not observed in control cells *in vivo*, we reasoned that our starting value of crosslinkers underestimated connectivity, potentially because cytoskeleton-membrane coupling is not accounted for in our minimal model. Therefore, we increased crosslinker abundance from our starting value and found that maximum ring closure speed peaked at 36,000, which suggested that cytoskeletal connectivity helps drive ring closure. This gain in ring speed was lost again by further increases in crosslinker abundance, indicating that they also attenuate ring closure.

**Figure 2.**
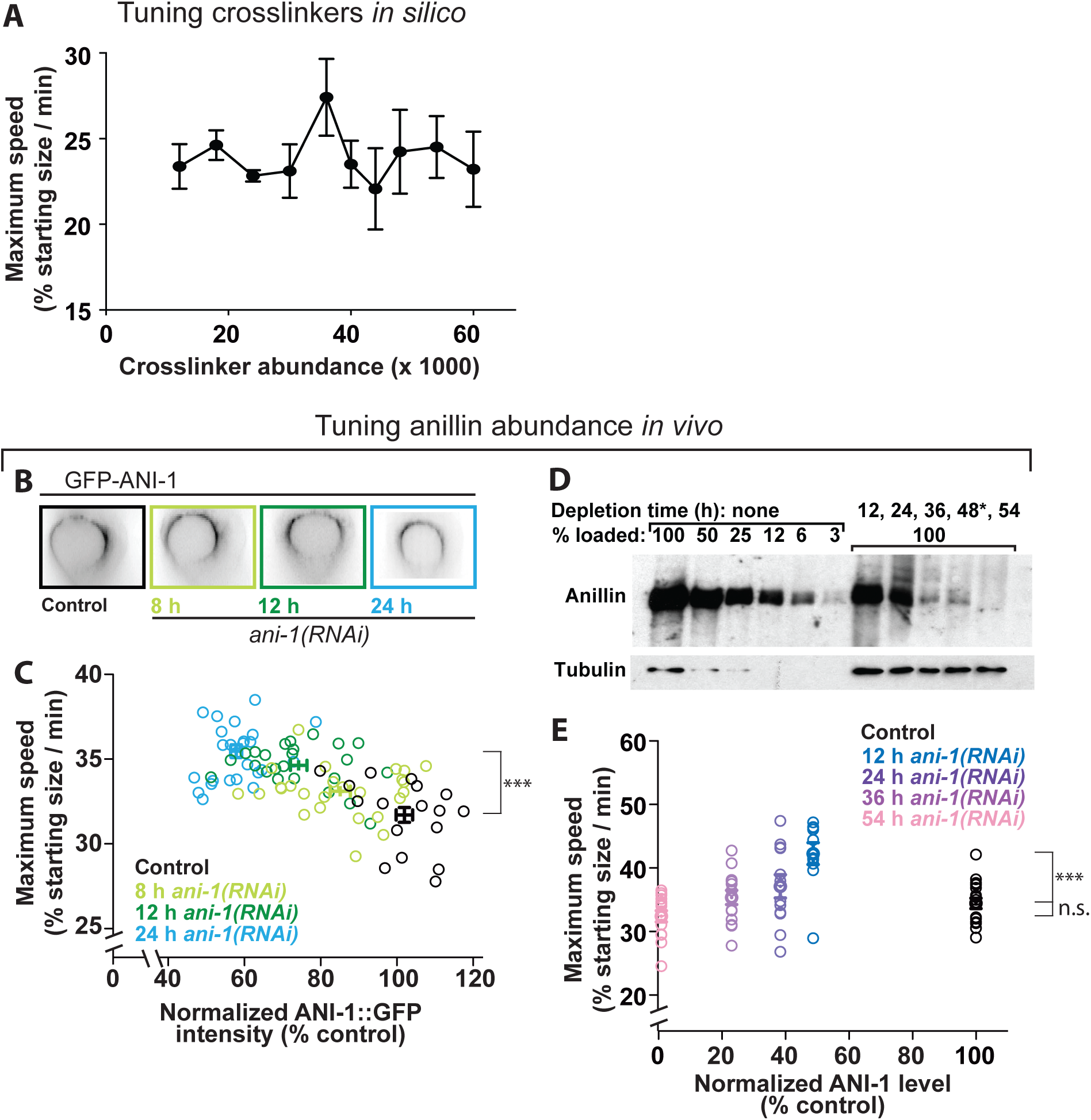
Non-motor crosslinkers both drive and brake cytokinetic ring closure in silico and in vivo. A) Maximum ring closure speed for simulations varying according to total non-motor crosslinkers (12,000 to 60,000). Averages ± SD for n = 4 simulations per condition are plotted. B) mNG::ANI-1 in the cytokinetic ring in representative control cell and those generated by worms depleted of anillin for varying amounts of time (hours). C) Maximum furrowing speed (at 50% closure) plotted against mNG::ANI-1 measured in the cytokinetic ring as in Figure 1E, for control cells and those depleted of anillin for between 8 and 24 h via dsRNA feeding. D) Western blots demonstrate the depletion of anillin over a time-course of injection RNAi. Tubulin = loading control. E) Maximum furrowing speed (at 50% closure) plotted against levels of residual endogenous anillin measured by western blotting, for control cells and those depleted of anillin for between 12 and 54 h via dsRNA injection. For C and E, bars denote average ± SEM.

We next tested the prediction that an intermediate level of non-motor crosslinkers *in vivo* also allows maximum speed. The multi-domain scaffold protein anillin is implicated in providing mechanical connectivity to the ring. While *Drosophila* and mammalian cells fail cytokinesis following anillin depletion, *C. elegans* zygotes are robust to thorough depletion of anillin (encoded by the gene *ani-1*; (Maddox *et al.*, 2005), allowing us to study the effects of depletion of cytokinetic dynamics. To test how the speed of cytokinetic ring constriction scales with the amount of crosslinking in animal cells, we depleted anillin over a time-course and measured residual anillin in the cytokinetic ring in embryos expressing anillin tagged at its endogenous locus with mNeonGreen (mNG::ANI-1; Figure 1D, 2B; (Rehain-Bell *et al.*, 2017)). The abundance of residual anillin in the cytokinetic ring was estimated from the fluorescence intensity in the 180 degree arc closest to the coverslip (Figure 1D). With increasing exposure time to bacterial food expressing dsRNA directed against *ani-1*, zygotes contained progressively less anillin (Figure 2B, C). We calculated the maximum furrowing speed (32 ± 0.6% of the starting radius per minute for control cells (Fig. 2C and C’; occurs at ~50% closure), as a metric for cytokinetic ring performance. As the level of anillin was reduced, furrowing reached a significantly higher maximum speed (p=0.0001; Figure 2C). These results suggest that, while anillin is a highly conserved crosslinker enriched in the cytokinetic ring and thus presumed to augment ring function, wild-type levels of anillin attenuate ring closure speed.

In cells depleted of anillin beyond approximately 50%, furrowing speed could no longer be reliably tracked by visualizing residual mNG::anillin. To test how more thorough anillin depletion changed furrowing speed, we tracked furrowing kinetics in worms expressing functional fluorescently-tagged NMM-II heavy chain (NMY-2::GFP) under the control of an integrated exogenous locus (Nance *et al.*, 2003). We measured residual endogenous anillin levels using western blots of whole worm extracts from worms depleted of anillin for various lengths of time, which led to a gradual reduction of anillin to 2% control levels (Figure 2F). Similar to our observation above, partial depletion of anillin to 50% of control levels lead to a significant increase in furrowing speed (p=0.0002; Figure 2E). Faster speeds at intermediate anillin levels could not be explained by an overabundance of NMM-II in the cytokinetic ring (Figure S1). In contrast, further reduction of anillin levels gradually restored furrowing speed to statistically indistinguishable from that of control cells (p=0.10 for 54 hour depletion compared to control; Figure 2G). These results suggest that, as *in vitro, in silico* (Ennomani *et al.*, 2016), and *in vivo* for plastin (Ding *et al.*, 2017), an intermediate amount of cytoskeletal crosslinking by anillin allows maximal contractility, and thus that crosslinking not only drives cytokinetic ring closure, but also attenuates furrowing speed. Furthermore, our *in silico* and *in vivo* results demonstrate that while alteration of crosslinker abundance can tune ring closure speed, these systems are robust, attaining control-level speed even after dramatic (50-fold) reduction of a major crosslinker. This is in contrast to the behavior of other simulated contractile networks, which fail to constrict with low crosslinker level and arrest with high crosslinker level (Ennomani *et al.*, 2016; Ding *et al.*, 2017). This distinction suggested that our simulated actomyosin network contained another robust crosslinking activity, namely our bipolar motors that represent NMM-II.

### NMM-II both drives cytokinetic ring closure and limits its maximum speed

The crosslinking activity of NMM-II has been implicated in driving cytokinetic ring closure (Ma *et al.*, 2012), but the effect of tuning motor-crosslinker levels on ingression dynamics has not been explored. To test if motor crosslinkers can both promote and attenuate ingression speed as described for non-motor crosslinkers, we first used our model to test how tuning the abundance of bipolar motor ensembles affected ring closure speed *in silico*, by performing simulations with randomly varied motor abundance between 20 and 120% of the control value (Figure 3A). Reduction of motor levels below 50% control slowed ring closure, in agreement with the idea that motors drive ring closure *in silico*. Furthermore, rings lacking bipolar motor arrays altogether failed to constrict (Figure 3B). However, over much of the range of motor level (between 50 and 120% control levels), ring closure speed did not appreciably vary (Figure 3A). The observation that adding more motors beyond a certain level does not increase ring closure speed supports the idea that bipolar motor arrays not only drive, but also attenuate, furrowing *in silico*.

**Figure 3.**
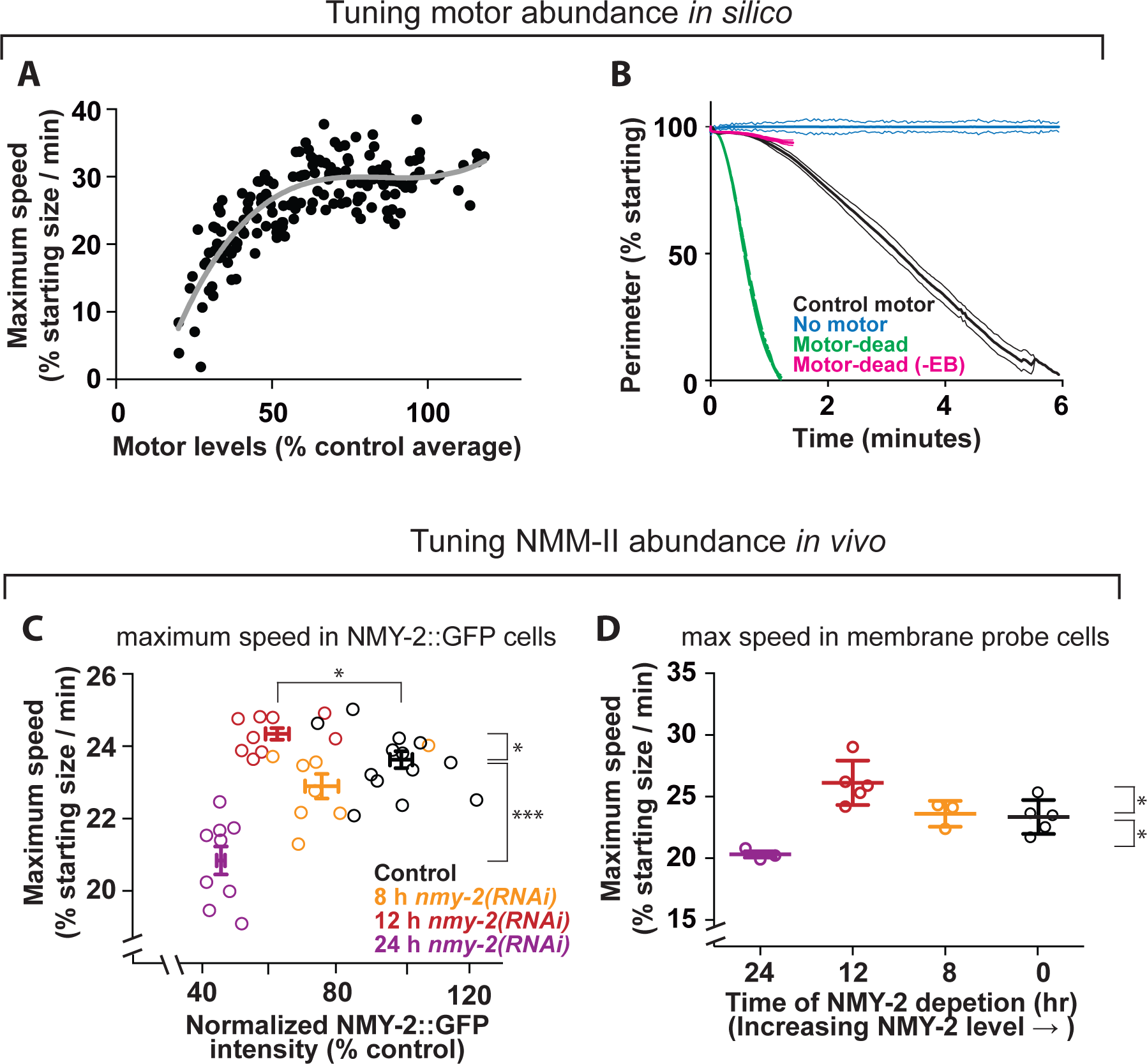
Furrowing kinetics respond non-linearly to NMM-II abundance. A) Maximum ring closure speed in simulations in which bipolar motor abundance was varied randomly between 20 and 120% control levels (n=167). Grey line: third-order polynomial (see Figure 4A and B). B) Simulated closure kinetics for rings containing control motor ensembles, motor-dead bipolar “motors” capable (“Motor dead”) or incapable of tracking fiber ends (“Motor dead (-EB)”), or lacking bipolar motor ensembles (“No motor”). Bars = SD; n=5 per condition. C) Maximum furrowing speed (at 50% closure) plotted against ring NMY-2::GFP for control cells and those depleted of NMY-2 for between 8, 12 and 24 h. D) Maximum furrowing speed (at 50% closure) in cells expressing a GFP-tagged membrane probe is compared for control cells and those depleted of NMY-2 for 8, 12 or 24 h. Two-tailed t-tests to compare control and 12 h: p=0.025; control versus 24 h: p=0.011.

To test how the level of NMM-II affects furrowing speed *in vivo*, we tuned ring levels of NMM-II by depleting the NMM-II heavy chain NMY-2 in worms expressing NMY-2-GFP, and measured residual NMY-2 to track ring closure. Reduction of NMY-2 levels below 50% of control levels significantly slowed ring closure (p<0.0001), in agreement with the idea that motors drive ring closure *in vivo*. However, partial depletion of NMY-2, to approximately 60% of control levels led to a significant increase in maximum ring closure speed (p=0.02), suggesting that NMY-2 can also attenuate closure speed *in vivo.* The same non-linear effect of myosin level on maximum furrowing speed was also obtained with a different *C. elegans* strain (Figure 3D). These results suggest that myosin not only drives ring closure but also resists cytoskeletal remodeling during furrowing.

Does NMM-II act as a motor, a crosslinker, or both? Some insight into this question may emerge from our minimal model. If NMM-II were to act only as a motor, one might expect that the speed of contractile events would scale linearly with NMM-II abundance. If NMM-II were to act only as a crosslinker, one would expect contractility speed to peak at intermediate NMM-II levels (Ennomani *et al.*, 2016; Li *et al.*, 2016). We reasoned that a combination of the two effects (motor – linear, and crosslinker - quadratic) would result in a complex relationship between NMMII and contractility. We sought mathematical evidence that NMM-II levels *in vivo* affect speed in a linear, quadratic, or more complex fashion (evidence that it acts as a motor, a crosslinker, or both), by fitting our data comparing NMM-II abundance and ring closure speed to polynomial curves of increasing order and monitoring the quality of the fit. Our *in silico* data were fit better by higher order polynomials than by simpler curves, indicating that motor abundance has a complex (non-linear) relationship with ring closure speed (Figure 4A, B). The same tests performed on *in vivo* data indicated that the dependence of ring closure speed on NMM-II abundance was also best fitted by higher order polynomial curves (Figure 4C and D). These findings support the idea that NMM-II functions both as a motor and a crosslinker.

**Figure 4.**
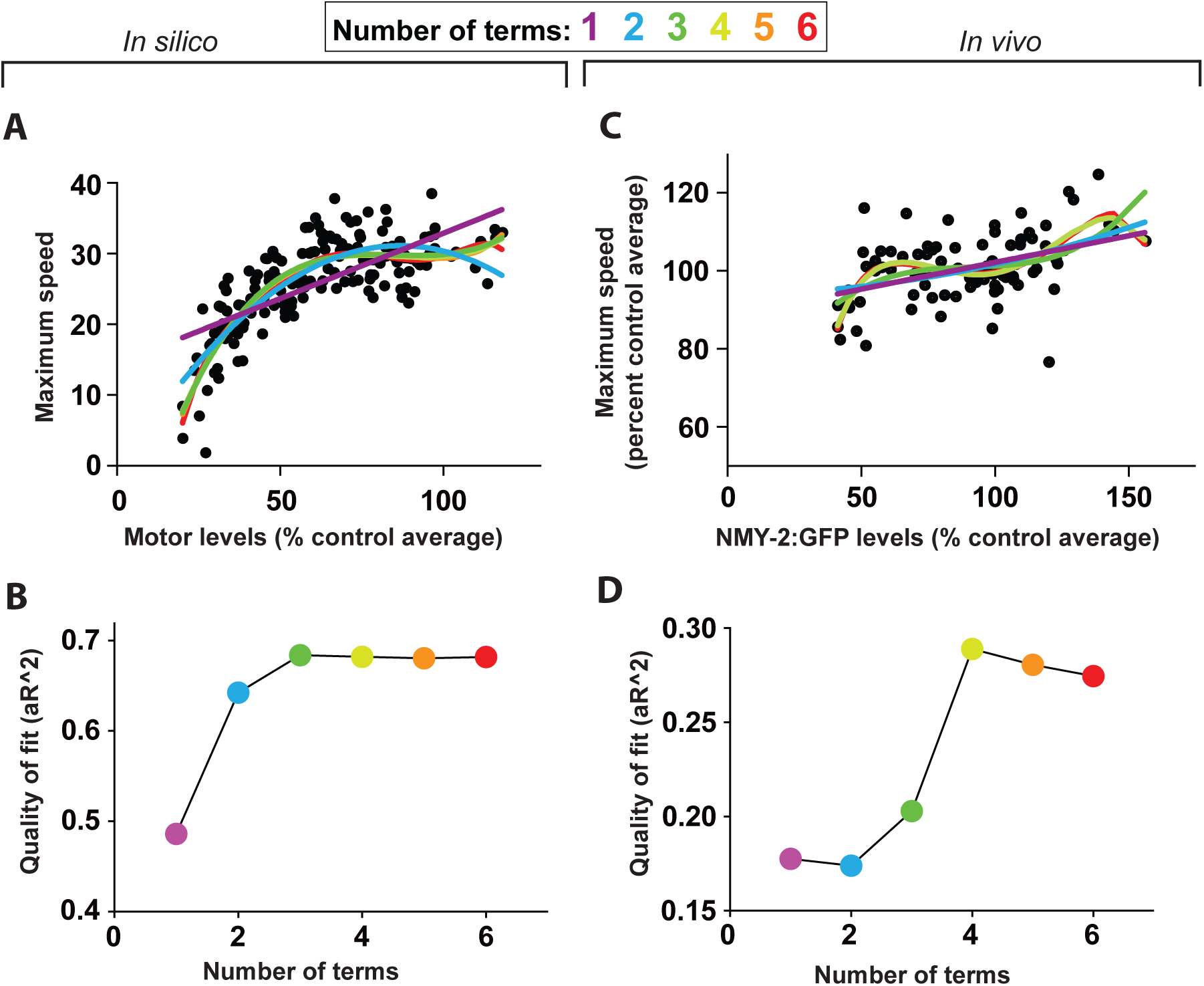
Motor abundance has a complex (non-linear) relationship with ring closure speed. A) Maximum ring closure speed for simulations in which bipolar motor abundance was varied randomly between 20 and 120% control levels (n=167) is duplicated from Figure 3A. B) Adjusted R^2^ values are plotted for each polynomial shown in A. C) Maximum ring closure speed data for cells in which NMM-II abundance varied via depletion of NMY-2 (data in Figure 3C) or the myosin-binding subunit of myosin phosphatase (Figure S3), normalized to the average of the appropriate control and plotted together with both sets of controls. Polynomial trendlines with increasing complexity were then fitted to the in vivo ring closure data. D) Adjusted R^2^ values are plotted for each polynomial in C.

Motor-impaired NMM-II is sufficient for cytokinesis in mammalian cells (Ma *et al.*, 2012). To recapitulate this result *in silico*, we simulated rings in which motor ensembles had a motoring speed of zero. Rings with “motor-dead” motor ensembles constricted faster than simulated ring with control motors (Figure 3B). This result agrees with the concept that crosslinking, in concert with treadmilling actin, can drive ring closure (Mendes Pinto *et al.*, 2012; Oelz *et al.*, 2015), and furthermore that motoring can oppose ring closure as a brake. Importantly, for motor-dead myosin-like ensembles to drive rapid ring closure *in silico*, they must track depolymerizing filament ends (Figure 3B). In sum, this *in silico* work supports the idea that the correct balance of motor and crosslinker activities and abundance is required for optimal ring dynamics.

### NMM-II both drives and attenuates cytokinetic furrowing initiation and F-actin organization

When the cytokinetic ring is at maximum speed, it is a tightly focused, diffraction limited cytoskeletal cord. In this configuration, crosslinkers could attenuate ring closure by blocking cytoskeletal sliding and remodeling. In contrast, when the cytokinetic ring first forms, cytoskeletal components are sparse, occupying a broad band of cortex (Henson *et al.*, 2017); in this configuration, crosslinkers could have different effects than at maximum speed. Thus, we next examined how NMM-II level relates to the timing of furrow initiation by again depleting NMMII over a time-course and scoring the onset of furrowing as time when the cortex first began to ingress away from the coverslip. Over most of the range of depletion time, initiation occurred later in cells more thoroughly depleted of NMM-II. However, when myosin was partially depleted to the level at which higher maximum speeds were achieved (12 hours), ingression occurred earlier than predicted by the trend exhibited by the other data (Figure 5A). These results suggest that while NMM-II drives furrow initiation, it also attenuates this early cell shape change to some degree.

**Figure 5.**
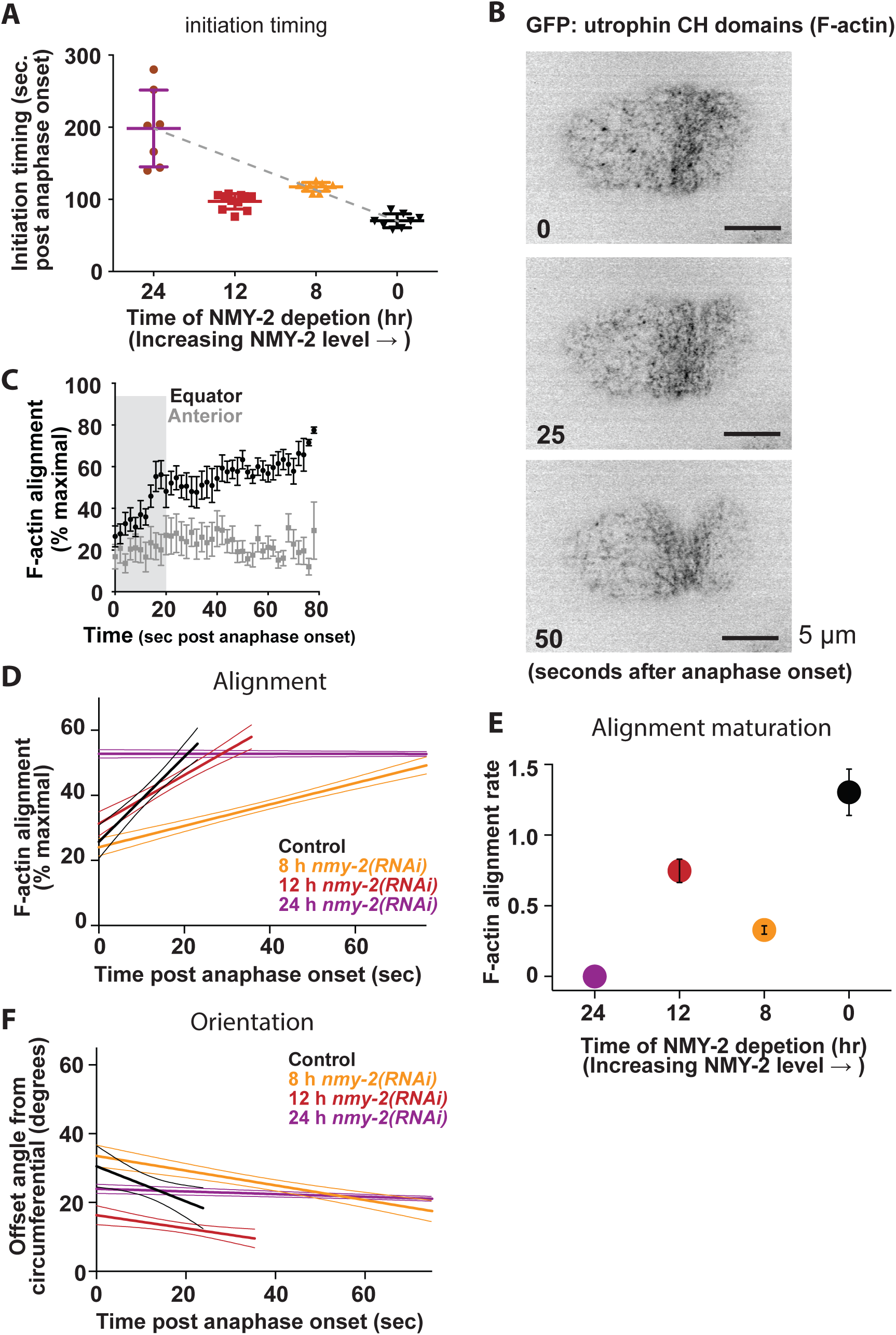
The kinetics of furrow initiation and circumferential F-actin bundling respond non-linearly to NMM-II abundance. A) Furrow initiation timing in cells expressing a GFP-tagged F-actin probe, relative to time of depletion for control cells and those depleted of NMY-2 for between 8 and 24 h. Bars denote average ± SEM. B) Cortical F-actin visualized via GFP-utrophin at three timepoints with respect to polar cap disappearance. C) F-actin order in the cell equator and anterior region (negative control); average ± SD are plotted; n =9. D) Initial linear increase in F-actin order in the cell equator plotted for control cells and those depleted of NMY-2 for 8-24 h. E) Average ± SD slope of initial linear increase in equatorial F-actin order for control cells and those depleted of NMY-2 for 8-24 h. For C, D and E, average ± SD are plotted; n > 7 cells per condition. F) Offset angle during the initial linear increase in F-actin order in the cell equator, plotted for control cells and those depleted of NMY-2 for 8-24 h. For C and D, thick lines = averages; thin lines = SD.

Furrow initiation has been attributed to the remodeling of an initially isotropic cytoskeletal meshwork into circumferential bundles (Reymann *et al.*, 2016). The effects of NMM-II levels on furrowing kinetics could relate to NMM-II’s roles in this cytoskeletal organization, but NMM-II could drive filament bundling and circumferential alignment, but also limit alignment by stabilizing isotropic filament intersections (Henson *et al.*, 2017). To test how these opposing activities affect early cytoskeletal remodeling during furrow initiation, we imaged cells expressing a GFP-tagged probe for F-actin (Jordan *et al.*, 2016), depleted NMY-2^MHC^ over a time-course, and measured the orientation of F-actin features, over time. As previously shown (Reymann *et al.*, 2016), F-actin is initially poorly organized in the cell equator, and local order increased in a linear fashion, in a process we will call F-actin alignment “maturation” (Figure 5B, C). The increase in order reflected a convergence on the circumferential direction (decrease in offset angle; see Figure 5F). We plotted the initial linear rate of maturation for control cells and those depleted of NMY-2 to varying extents. The rate of alignment maturation was reduced by the mildest depletion, and more strongly reduced by the most severe depletion (Figure 5D, E). However, at the intermediate NMM-II level at which higher maximum furrowing speed and precocious initiation took place, the rate of F-actin alignment maturation was higher than expected based on the maturation rates of control cells and those depleted for 8 or 24 hours. From these results, we conclude that NMM-II not only drives but also attenuates cytoskeletal alignment. This implicates NMM-II not only in crosslinking, but specifically in promoting filament alignment, potentially by mediating filament bundling. In sum, our findings support the idea that crosslinkers not only contribute positively to, but also limit, the cytoskeletal remodeling that underlies cytokinetic ring closure.

## Discussion

Speed is of the essence for cytokinesis in early embryos, since cell cycles are short, and a persistent cytoplasmic connection between daughter cells could lead to intercellular aneuploidy or defects in cell fate determination. Like other aspects of cell division in the *C. elegans* zygote, furrowing is exceptionally rapid and appears to have been fully optimized by evolutionary adaptation for high-fidelity, rapid early development. Therefore, it is somewhat unexpected that any experimental perturbation can increase furrowing speed. How can a depleted cytokinetic ring function “better” than a control ring? Crosslinkers, including NMM-II, may block cytoskeletal remodeling and therefore furrowing in one or more ways, including stiffening the cortex by stabilizing interactions, suppressing actin depolymerization, or via steric hindrance. One might predict that crosslinkers in excess of the levels that allow maximal furrowing speed are incorporated into the ring where they would facilitate early remodeling. However, the same intermediate NMM-II depletion that allowed faster maximum speed also resulted in faster-than-expected cytoskeletal organization and furrow initiation (Figures 3C and D, and 5E). This lack of kinetic advantage in any of the stages we assayed demonstrates that mechanical brakes are part of the cytokinetic ring despite compromising speed, and suggests that they offer a distinct advantage, such as to make furrowing responsive to inhibitory signals for error correction.

The nature of the cytoskeletal remodeling that drives animal cytokinetic ring closure is still debated. One hypothesis is that filaments slide past one another as in muscle contraction (Schroeder, 1972). Alternatively, the new ends of depolymerizing F-actin may be ratcheted together via dynamic non-motor crosslinking (Mendes Pinto *et al.*, 2012). Our findings that crosslinkers both drive and attenuate furrowing do not refute this latter model, but suggest that the abundance of crosslinker must be delicately balanced to optimally achieve remodeling, and thus extend our understanding of the structural mechanisms of cytokinetic ring closure.

Of similar importance is the question of how myosin motors drive constriction. Recent work has demonstrated that dynamic end-tracking crosslinking can drive constriction (Mendes Pinto *et al.*, 2012; Oelz *et al.*, 2015), but a role for motoring activity has not been fully discounted (Zumdieck *et al.*, 2007; Oelz *et al.*, 2015; Bidone *et al.*, 2017). Our *in silico* findings that motor-dead myosin-like ensembles can drive ring closure as long as they can track depolymerizing actin fiber ends (Figure 3B) support the idea that dynamic crosslinking of treadmilling actin drives contraction. However, we also found that motoring activity slows closure of *in silico* rings, compared to motor-dead myosins, supporting the idea that motoring attenuates actomyosin contractile dynamics. In sum, our work suggests three contributions from motor ensembles: driving contractility via motoring and via crosslinking, as well as attenuating contractility. These in silico findings must be considered cautiously, since motor-dead NMM-II *in vivo* could have different biophysical properties than *in silico*; it may not only lack the power stroke but also bind and unbind F-actin with different kinetics. In addition, our modeling of NMM-II minifilaments may not accurately simulate some aspects of motor ensembles; our calculated multimerization of binding and unbinding dynamics of NMMII minifilaments could change the effect or magnitude of motoring activity on contractility.

We found that an intermediate amount of anillin and NMM-II allows maximal contraction speed. Similarly, maximal speed results from intermediate amounts of alpha-actinin in *S. pombe* (Li *et al.*, 2016) and *in vitro* (Ennomani *et al.*, 2016). Agent-based modeling of contractile networks containing generic motor dimers (Ding *et al.*, 2017), myosin VI-like dimers (Ennomani *et al.*, 2016), bipolar motor ensembles that represent NMM-II minifilaments (Figure 1D) also demonstrates that intermediate amounts of crosslinking allows maximal contractility. Continuum theory suggests that this behavior results from an effect on cytoskeletal mesh size and structure (Lenz *et al.*, 2012). Together, these diverse observations suggest that the balance of positive and negative effects of motor and non-motor cytoskeletal crosslinkers, revealed here for animal cell cytokinesis, is a general principle of actomyosin contractility.

## Experimental procedures

### Strains, growth conditions and sample preparation

The following strains were used: JJ1473 *(zuIs45 [nmy-2*::*NMY-2*:: *GFP + unc-119(+)] V)*, OD95 *(unc-119(ed3) III; ltIs38 [pAA1; pie-1/GFP*::*PH(PLC1delta1); unc-119 (+)]; ltIs37 [pAA64; pie-1/mCherry*:: *his-58; unc-119 (+)] IV)*, MDX29 (ani-1(mon7[mNeonGreen^3xFlag::ani-1]) III) (Rehain-Bell *et al.*, 2017), MDX40 (ani-1(mon7[mNeonGreen^3xFlag::ani-1], unc-119(ed3) III; nmy-2(cp52[nmy-2::mkate2 + LoxP unc-119(+) LoxP]) I), and JCC719 (mgSi3[tb-unc-119(+) pie-1>gfp:utrophin] II.; unc-119 (+)] III; ltIs37 [pAA64; pie-1/mCherry::his-58; unc-119 (+)] IV) (Jordan *et al.*, 2016). Worms were maintained at 20°C according to standard procedures (Munro *et al.*, 2004). Embryos were dissected from gravid hermaphrodites and mounted on coverslips on 2% agarose pads.

### RNA-mediated protein depletions

Depletion of ANI-1, NMY-2, LET-502, and MEL-11 was performed by feeding worms bacteria from the Ahringer collection expressing dsRNAs directed against the targets, according to standard procedures (Min *et al.*, 2010). The identity and sequence of all dsRNAs were confirmed by sequencing. Throughout this study, we exclusively examined perturbations that allow cytokinesis to succeed, while exhibiting quantitative defects.

### *C. elegans* embryo live cell imaging conditions

Imaging for panels 2C-E, 3C-D, 4A, 5A and S2 was performed on a DeltaVision Elite widefield microscopy system using the software Softworks (Applied Precision GE-Healthcare), equipped with a CoolSnap HQ2 camera (Photometrics) at 2x2 binning and a 60x/1.42 plan Apo objective (Olympus). All image analysis was performed on non-deconvolved data.

Imaging for panels 2G and S1 (NMY-2::GFP embryos depleted of ANI-1^anillin^) was performed with a Nikon TE-2000 inverted microscope equipped with a swept field real-time confocal module (Prairie Instruments) and Coolsnap HQ2 CCD camera (Photometrics). The use of the 70-μm slit, 60x magnification, and a 200 ms exposure time were kept constant.

Imaging of actin alignment during furrow initiation was performed on a Nikon TE300 inverted microscope with a Plan APO 60x 1.4 NA oil objective, a Yokogawa Nipkow spinning disk, Hamamatsu ORCA-ER camera, Prior stage microcontroller, and Melles Griot ion laser power supply. Embryos were dissected, as previously described, and imaged prior to 1 cell division with 2 second intervals between each image acquisition. Image data was aligned based on disappearance of the polar concentration of actin and the onset of cortical swirling prior to furrow initiation. Images were acquired in a single cortical plane through the maturation of the furrow, which was determined based on the ingression of the contractile ring past the cortical imaging plane.

### Measurements of ring closure

Anaphase onset was determined by observing a separation between chromatin masses visible against the background of myosin fluorescence in the spindle. Ring closure was determined using semi-automated custom software cyanRing (for CYtokinesis ANalysis of the RING) as described previously (Dorn *et al.*, 2010; Bourdages *et al.*, 2014). In cyanRing, the equatorial region of the embryo is cropped from the 3D image stack, rotated, and maximum-intensity projected to produce a view of the cytokinetic ring along the spindle axis as viewed from the posterior direction. The user then defines the cell outline as well as the outline of the cytokinetic ring by selecting at least three points, through which best-fit circles are drawn. Radii are calculated from best-fit circles and used to calculate ring closure as a percentage of starting size.

### Measurements of cytokinetic ring myosin levels

The intensity of myosin at maximum speed was extracted from cyanRing annotations of ring position. The integrated intensity was calculated for a 9-pixel radius torus encompassing the cytokinetic ring and normalized against a region of background outside the cell. Experimental intensity values were normalized to the average of control cell data acquired on the same day.

### Quantitative immunoblotting

L4-stage hermaphrodites were injected with dsRNA directed against *ani-1* (Maddox et al., 2005) and incubated at 20°C for 12-54 hours. Extracts were prepared from control worms and those depleted of ANI-1^anillin^ using standard protocols. Extract loaded onto the 100% loading lanes contains approximately 10 worms. Immunoblotting was performed using an anti-ANI-1 polyclonal antibody (Maddox et al., 2005) and anti-α-tubulin clone DM1α (Sigma). Protein bands from film were quantified using ImageJ software. Loading was normalized using tubulin as a control. Data presented are the relative anillin amounts as a ratio of anillin in depleted worms relative to that of control worms. The extent of depletion estimated via blotting for endogenous anillin in whole worms likely underestimates levels in embryos, since the target protein is more thoroughly depleted from the germline and embryos than other tissues.

### Simulations

Agent-based modeling was performed using the Open Source software CytoSim beta for MacOS 10.6+ ((Nedelec and Foethke, 2007), <u>www.cytosim.org</u>). General setup is described in great detail in the included simulation file and supplemental table (Supplemental File 1; Supplemental Table 1). Motor head unbinding dynamics are calculated by the Kramer method: k_off_ = k_0_ * exp(F_t_ / F_0_); where k_0_ is the unloaded unbinding rate, F_0_ is the unbinding force, and F_t_ is the force felt by the binding motor at time t. Motoring speed was calculated at each simulation frame by the formula: V_t_ = V_0_ * (1-F_t_ / F_0_); where V_0_ is the unloaded motor speed, F_0_ is the stall force, and F_t_ is the load force at timepoint t. Myosin complexes were simulated as bipolar couples with actin-binding motoring domains attached at either end of a filament. Each actin-binding motor domain was modeled to simulate the dynamics of multiple myosin motoring heads (between 2-28) for total myosin valence of 4-56 motors. Actin dynamics were based off previous models (Mendes Pinto *et al.*, 2012; Oelz *et al.*, 2015) and include depolymerizing primarily at pointed ends and polymerizing primarily at barbed ends, with a net treadmilling rate of 0.04 μm/s and net depolymerization rate of 0.004 μm/s. Crosslinkers were modeled after those of previous work (Ennomani *et al.*, 2016) to represent a crosslinker like alpha actinin, bearing a single actin-binding domain at each end of a small filament. Initially, actin filaments were assembled in a tangential orientation to the circular cell space in a radius of approximately 14.5 μmwith a ring width of 300 nm. After equilibrating in the simulation crosslinkers and motors were added to the simulation space and simuations were allowed to run for 400 seconds of simulated time. Control contractile ring simulation parameters are listed in supplemental table 1 and in the attached configuration file. Plots of *in silico* ring closure kinetics were smoothened with a five time-point rolling average to reduce noise.

### Stepwise regression analysis

Stepwise regression analysis for our *in vivo* and *in silico* myosin depletion speed curves was performed using GraphPad Prism software (GraphPad Software Inc, La Jolla, CA). Data were normalized to control myosin levels (x axis) and control maximum speed (y axis). Polynomial regressions were fitted to the normalized experimental data and simulation data, and F-test analysis was performed to generate absolute sum of squares and adjusted R^2^ (aR^2^) values for each polynomial fit. Optimal fitting was determined based on increase of the aR^2^ to a peak value followed by plateauing or decrease as additional terms are added to the polynomial fit equation.

## Acknowledgements

We are grateful to Laura Miller, Vincent Boudreau, Young Jun Yun, Xiaohu Wan, Kathryn Rehain-Bell, and Melissa Plooster for technical help. We thank Jian Liu for insightful discussions and Mark Peifer, Michael Werner, and Dan Dickinson for critical reading of this manuscript. We thank the entire Maddox labs for their thoughtful input, and Scott Williams for his support.

## Supplemental File 1

The CytoSim configuration file is provided as a supplemental document.

## Supplemental Table 1

| Component | Properties | Comments |
| --- | --- | --- |
| Simulation | Space: circle, 15 $\mu\text{m}$ radius<br>Viscosity: 1 N.s.m<sup>2</sup><br>Temperature: 0.0042 pN.um<br>Time-step: 2 ms | Viscosity corresponds to typical measurement for embryo cytoplasm (Daniels *et al.*, 2006) |
| Actin-like fibers | 2400 fibers<br>Length: 1.3 ± 0.3 $\mu\text{m}$<br>Rigidity: 0.06 pN.$\mu\text{m}^2$<br>Barbed end growth speed: 0.054 $\mu\text{m/s}$<br>Pointed end growth speed: 0.01 $\mu\text{m/s}$<br>Barbed end shrinking speed: 0.01 $\mu\text{m/s}$<br>Pointed end shrinking speed: 0.05 $\mu\text{m/s}$ | Filaments treadmill |
| Motor head | Binding rate: 0.2 /s<br>Binding range: 330 nm<br>Unbinding rate: 0.3/s<br>Unbinding force: 3.85 pN<br>Max speed: 100 nm/s<br>Stall force: 5 pN<br>Speed formula: $V=V_0 * (1-F/F_0)$ | All parameters are per motor head<br>For ensemble stall force and unbinding force multiply by (valence * duty ratio) |
| Motor ensembles | 1800 ensembles<br>Valence: 14 per “hand”<br>Length: 320 nm<br>Stiffness: 1400 pN/$\mu\text{m}$<br>Duty ratio: 0.35<br>Diffusion: Instantaneous | Ensembles track depolymerizing actin pointed ends<br>Duty ratio is assumed to be invariant for the whole minifilament ensemble |
| Crosslinkers | 44000<br>Length: 40 nm<br>Binding rate: 10/s<br>Binding range: 100 nm<br>Unbinding rate: 0 /s<br>Unbinding force: 5 pN<br>Stiffness: 100 pN/$\mu\text{m}$<br>Diffusion: Instantaneous | Crosslinkers are removed at a rate of 2400/ 25s for 50 sec, then at a rate of 2850/ 25s |

**Supplemental Figure 1.**
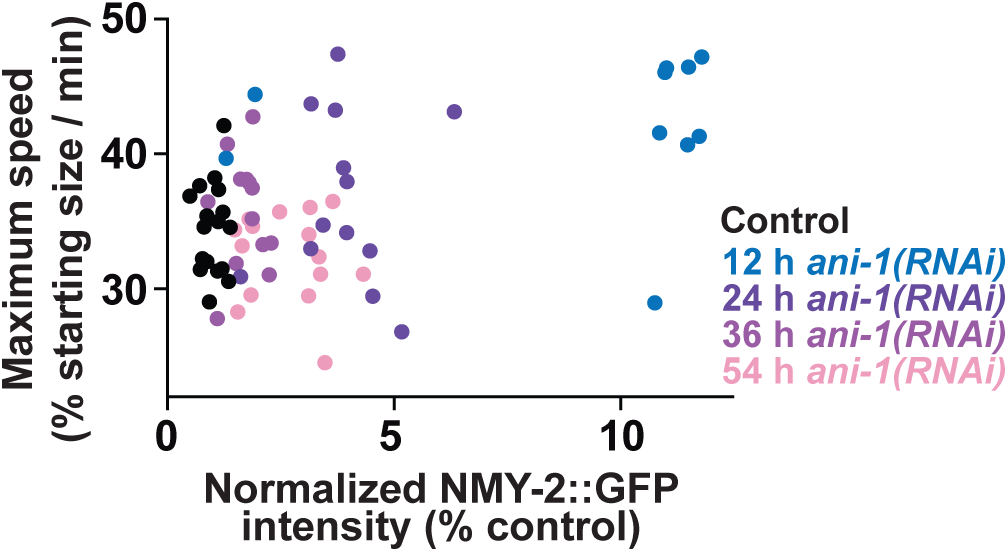
(Relating to Main Figure 2G) Furrowing speed does not correlate with NMY-2::GFP levels in cells depleted of anillin.

**Supplemental Figure 2.**
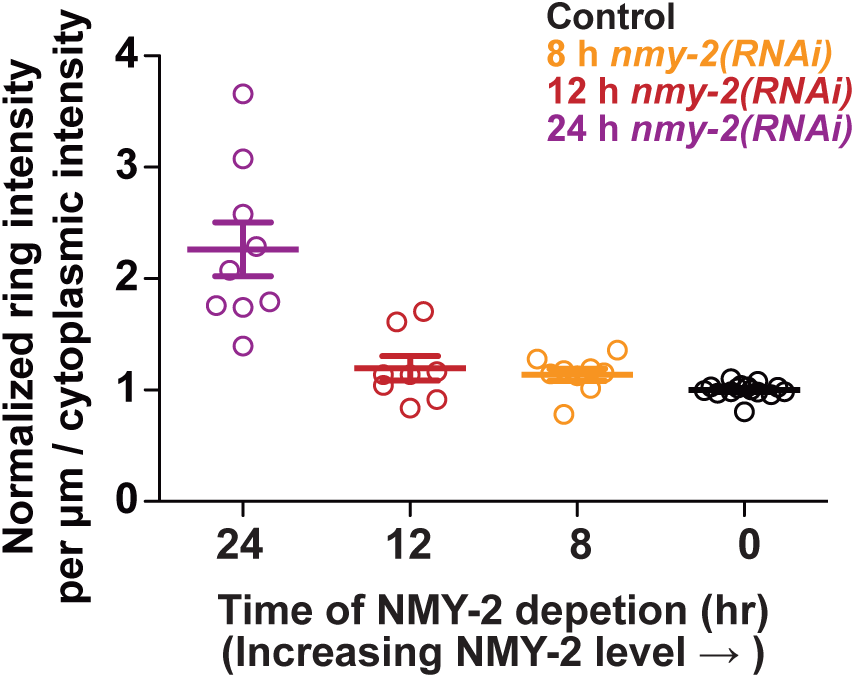
(Relating to Main Figure 3) Only a relatively thorough partial depletion of NMY- 2 affects the enrichment of NMY-2::GFP on the cytokinetic ring compared to cytoplasmic levels. Ring enrichment was calculated as the fluorescence intensity of NMY-2::GFP on the ring relative to cytoplasmic background (see Figure 2A). Values are normalized to average of control cell value. Bars = average + SEM.

**Supplemental Figure 3.**
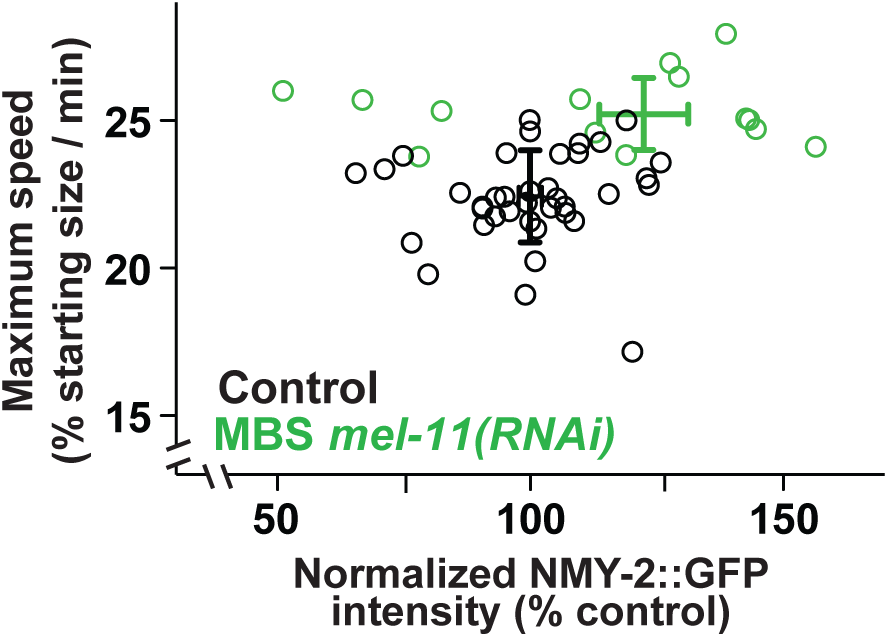
(Relating to Main Figure 4C) Furrowing speed at 50% closure plotted against NMY-2::GFP measured in the cytokinetic ring as in Fig 2A, for control cells and those depleted of the myosin-binding subunit of the myosin phosphatase (MEL-11).

**Supplemental Figure 4.**
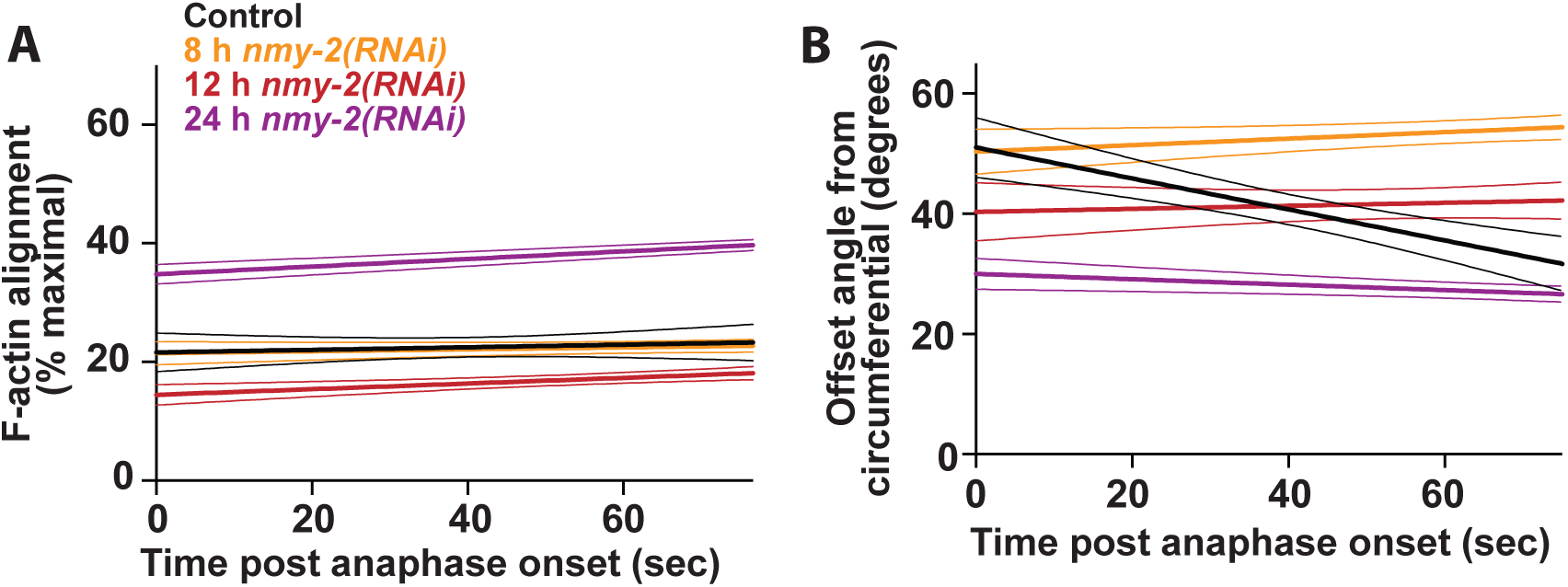
(Relating to Main Figure 5) Cortical F-actin in the cell anterior did not exhibit maturation of circumferential alignment. A) change in order over time. B) change in dominant angle, with respect to circumferential (0 degrees).

